# Differential expression of RET and GDNF family receptor, GFR-α1, between striatum and substantia nigra following nigrostriatal lesion: a case for diminished GDNF-signaling

**DOI:** 10.1101/2023.03.01.530671

**Authors:** Ella A. Kasanga, Yoonhee Han, Walter Navarrete, Robert McManus, Marla K. Shifflet, Caleb Parry, Arturo Barahona, Fredric P. Manfredsson, Vicki A. Nejtek, Jason R. Richardson, Michael F. Salvatore

## Abstract

Although glial cell line-derived neurotrophic factor (GDNF) showed efficacy in preclinical and early clinical studies to alleviate parkinsonian signs in Parkinson’s disease (PD), later trials did not meet primary endpoints, giving pause to consider further investigation. While GDNF dose and delivery methods may have contributed to diminished efficacy, one crucial aspect of these clinical studies is that GDNF treatment across all studies began ∼8 years after PD diagnosis; a time point representing several years after near 100% depletion of nigrostriatal dopamine markers in striatum and at least 50% in substantia nigra (SN), and is later than the timing of GDNF treatment in preclinical studies. With nigrostriatal terminal loss exceeding 70% at PD diagnosis, we utilized hemi-parkinsonian rats to determine if expression of GDNF family receptor, GFR-α1, and receptor tyrosine kinase, RET, differed between striatum and SN at 1 and 4 weeks following a 6-hydroxydopamine (6-OHDA) lesion. Whereas GDNF expression changed minimally, GFR-α1 expression decreased progressively in striatum and in tyrosine hydroxylase positive (TH+) cells in SN, correlating with reduced TH cell number. However, in nigral astrocytes, GFR-α1 expression increased. RET expression decreased maximally in striatum by 1 week, whereas in the SN, a transient bilateral increase occurred that returned to control levels by 4 weeks. Expression of brain-derived neurotrophic factor (BDNF) or its receptor, TrkB, were unchanged throughout lesion progression. Together, these results reveal that differential GFR-α1 and RET expression between the striatum and SN, and cell-specific differences in GFR-α1 expression in SN, occur during nigrostriatal neuron loss. Targeting loss of GDNF receptors appears critical to enhance GDNF therapeutic efficacy against nigrostriatal neuron loss.

**Significance Statement:** Although preclinical evidence supports that GDNF provides neuroprotection and improves locomotor function in preclinical studies, clinical data supporting its efficacy to alleviate motor impairment in Parkinson’s disease patients remains uncertain. Using the established 6-OHDA hemi-parkinsonian rat model, we determined whether expression of its cognate receptors, GFR-α1 and RET, were differentially affected between striatum and substantia nigra in a timeline study. In striatum, there was early and significant loss of RET, but a gradual, progressive loss of GFR-α1. In contrast, RET transiently increased in lesioned substantia nigra, but GFR-α1 progressively decreased only in nigrostriatal neurons and correlated with TH cell loss. Our results indicate that direct availability of GFR-α1 may be a critical element that determines GDNF efficacy following striatal delivery.

**Highlights:** GDNF expression was minimally affected by nigrostriatal lesion

GDNF family receptor, GFR-α1, progressively decreased in striatum and in TH neurons in SN.

GFR-α1 expression decreased along with TH neurons as lesion progressed

GFR-α1 increased bilaterally in GFAP+ cells suggesting an inherent response to offset TH neuron loss

RET expression was severely reduced in striatum, whereas it increased in SN early after lesion induction

## Introduction

Identifying disease-modifying therapies to slow nigrostriatal neuron loss is an unmet need in Parkinson’s disease (PD) therapy (AlDakheel et al., 2014; Barker et al., 2020; Björklund et al., 2021; Manfredsson et al., 2020; Paul and Sullivan, 2019). This challenge may be further compounded by the complexities of PD etiopathology (Kalia and Lang, 2015), detecting PD pathology prior to onset of motor impairment (Berg et al., 2021; Blesa et al., 2022; Heinzel et al., 2019; Salvatore et al., 2021), and still limited knowledge of the mechanisms underlying the neurodegenerative process in nigrostriatal neurons (Chmielarz and Saarma, 2020; Espay et al., 2020; Kalia et al., 2015; Zeng et al., 2018).

Neurotrophic factors such as brain-derived neurotrophic factor (BDNF) and glial cell line-derived neurotrophic factor (GDNF) have been investigated as potential neurodegenerative therapies (Airaksinen and Saarma, 2003; Burke, 2006; Chmielarz and Saarma, 2020; Ferreira et al., 2018; Palasz et al., 2020; Tome et al., 2017). This is primarily due to their ability to support the survival of neurons and increase function and connectivity (Chmielarz and Saarma, 2020). BDNF promotes the survival of dopaminergic neurons in animal models of PD, however, unlike GDNF, BDNF efficacy may be limited in early-stages of nigrostriatal neuron loss PD (Chmielarz and Saarma, 2020). GDNF on the other hand, has been widely investigated (Gash et al., 1996; Grondin et al., 2018; Kasanga et al., 2019; Manfredsson et al., 2020; Salvatore et al., 2009, 2004) since its discovery in the early 90s (Lin et al., 1993). Unfortunately, clinical trials, so far, have produced inconsistent results. Whereas initial phase I trials achieved their primary outcomes, with improvements in UPDRS scores (Gill et al., 2003; Slevin et al., 2005), subsequent trials did not meet primary endpoints (Lang et al., 2006; Whone et al., 2019). Some of the reasons for failure in later trials include insufficient GDNF dose, duration of delivery, and delivery method of GDNF (Ai et al., 2003; Bartus and Johnson, 2017; Gash et al., 2005; Kasanga et al., 2019; Luz et al., 2018; Salvatore et al., 2006; Sherer et al., 2006). A common theme for all trials was that degree of nigrostriatal terminal loss at the time of GDNF delivery was likely beyond complete. The mean time of GDNF delivery from PD diagnosis was ∼8 years, and terminal loss is virtually complete by 5 years diagnosis (Kordower et al., 2013). Thus, the responsivity of nigrostriatal terminals to GDNF, namely via the availability of GDNF receptors, at that degree of loss could be severely compromised and reduce potential for plasticity in dopamine signaling in residual nigrostriatal neurons (Chu and Kordower, 2021; Espay et al., 2020; Quintino et al., 2019).

GDNF is a potent dopaminergic neurotrophic factor which is normally expressed at low levels in the adult brain (Liberatore et al., 1997; Strömberg et al., 1993). However, the role of the GDNF family receptor, GFR-α1, in mediating neuroprotection in PD has been sparsely studied. Global knockout of GFR-α1 leads to death of mice soon after birth (Enomoto et al., 1998), however, heterozygous GFR-α1 knockout mice develop decreased motor function concurrent with a reduction in nigral dopaminergic neurons with aging (Zaman et al., 2008). RET, the GDNF/GFR-α1 activated receptor tyrosine kinase, has been more widely investigated for a role in nigrostriatal function. Small molecules targeting RET receptors stimulate neuronal signaling and dopamine release indicating their therapeutic potential (Conway and Kramer, 2021; Kramer and Liss, 2015; Mahato and Sidorova, 2020). Post-mortem analysis of putamen samples from PD subjects revealed no significant changes in GFR-α1 and RET mRNA expression with a modest increase in GDNF mRNA levels and significant loss of nigral neurons (Bäckman et al., 2006). In the SN, decreased RET expression suggests a key role in nigrostriatal neuron viability (Chu and Kordower, 2021). Moreover, RET activation differentially protects dopaminergic cell bodies instead of axon terminals after neurotoxic insult, indicating RET signaling in the SN is critical for neuroprotection (Mijatovic et al., 2011).

Given the physiological impact of these receptors on nigrostriatal function, we evaluated the short and long-term impact of nigrostriatal neuron loss on expression of GDNF and BDNF, and cognate receptors, after 6-hydroxydopamine (6-OHDA) lesion in rats. Comprehensive analysis of relative differences in expression of these receptors in nigrostriatal terminals vs cell bodies revealed critical time windows and nigrostriatal compartments of when and where in the nigrostriatal pathway GDNF efficacy to recover or protect DA function in the nigrostriatal pathway is feasible or limited.

## Methods

### Animals

Male Sprague-Dawley rats (3-month-old, n=65) were purchased from Charles River (Worcester, MA, USA). Rats were housed under controlled lighting conditions with standard animal chow and water available *ad libitum*. Animals were used in compliance with the National Research Council’s Guide for the Care and Use of Laboratory Animals and protocols (IACUC-2021-0018) approved by the Institutional Animal Care and Use Committee at the University of North Texas Health Science Center.

### Experimental Design

Two cohorts were employed in this study – a 7-day (n=33) and 28-day cohort (n=32). Each cohort consisted of a sham and 6-OHDA group. Rats underwent unilateral nigrostriatal lesioning, as described below, after which rats were euthanized at either 7- or 28-days post-lesioning after locomotor testing using the forepaw adjusting steps (FAS) task. Rats were further divided into a neurochemistry cohort (n=40) for the analysis of GDNF, BDNF, and their receptors via quantitative western blot and an immunohistochemistry cohort (n=25) for the analysis of GFR-α1 and RET counts. The impact of nigrostriatal lesioning on locomotor impairment and DA regulation in the striatum vs SN with this experimental design, and reflecting these measures in the rats used in this study, has been published (Kasanga et al., 2022).

### 6-OHDA nigrostriatal lesion induction

Rats were anesthetized with 2-3% continuous inhalation isoflurane for survival surgery to deliver 6-OHDA (Chotibut et al., 2017, 2014). Briefly, rats were immobilized in a stereotaxic frame to target the medial forebrain bundle (mfb) at coordinates relative to Bregma (ML - 2.0, AP - 1.8, DV - 8.6) (Paxinos and Watson, 2014). Rats either received 6-OHDA (16µg in 4µL of vehicle (0.02%^w^/_v_ ascorbic acid) in the left mfb or vehicle. In the lesioned group only, the vehicle was infused into the contralateral mfb. The needle was left in place for 10 min before removal. Body temperature was maintained at 37°C using a temperature monitor and heating pad. Nigrostriatal lesioning was confirmed by FAS at day 7 for the neurochemistry cohort (Kasanga et al., 2022). In the immunohistochemistry cohort, striatal tyrosine hydroxylase (TH) loss ≥80% in lesioned striatum versus levels in contralateral striatum was used as the criterion for a successful nigrostriatal lesion. A total of 3 rats (out of 14) did not meet this inclusion criterion.

### Forepaw Adjusting Steps Task

FAS is used to measure forelimb akinesia resulting from DA depletion (Chotibut et al., 2017; Meadows et al., 2017). Briefly, both hind limbs and one forepaw are held such that the rat bears its weight on the forepaw to be tested. The rat is then moved across a table at a speed of 90cm/10s, during which the number of adjusting steps taken by the rat is counted. This is done for 6 trials per forepaw: 3 forehand trials (lateral steps toward the thumb of the paw that is stepping) and 3 backhand trials (lateral steps towards the pinky of the paw that is stepping), alternating the starting forepaw between rats. FAS revealed a significant reduction (≥75%) in contralateral forepaw use (to the lesioned hemisphere) as compared to the ipsilateral forepaw at day 7 in both the 7-day cohort (t=7.41, *p* <0.0001, df=8) and 28-day cohort (t=11.77, *p* <0.0001, df=8) with no significant difference between the two cohorts (data not shown). Rats with <25% contralateral forelimb use (6 out of 25) were excluded from further analysis (Kasanga et al., 2022).

### Tissue Processing

Brains for neurochemical analyses were processed as previously described (Salvatore et al., 2012; Salvatore and Pruett, 2012). Briefly, rats were euthanized at the respective time points with tissue punches of striatum, or hand-dissected SN collected on wet ice, immediately cooled on dry ice and saved for subsequent western blot processing.

Tissue punches of the striatum were collected on wet ice from samples allocated for immunohistochemical analysis (n=25) to verify lesion efficacy. The portion containing the midbrain sections were post-fixed in 4% paraformaldehyde for 7 days at 4℃ after which the tissue was immersed in a 30% sucrose + 0.1% sodium azide solution at 4℃ for cryoprotection until sectioning. Immunohistochemistry was performed on free-floating sliding microtome-cut sections (30μm in thickness) encompassing the SN.

### Determination of Protein Expression

SDS gel electrophoresis was conducted after total protein quantitation assay. Following protein transfer, the nitrocellulose membranes were stained with ponceau S and imaged for subsequent normalization to total protein, blocked for at least 2 hours, and subsequently placed in the respective primary antibody solutions at 4℃ overnight. The specific antibodies used include TH (anti-rabbit-TH; AB152; 1:1000; Millipore, Temecula, CA), GDNF (anti-goat-GDNF; AF-212-NA, 0.2µg/ml, R&D Systems, Minneapolis, MN), GFR-α1 (anti-goat-GFR-α1; AF560; 1µg/ml; R&D Systems, Minneapolis, MN), RET (anti-rabbit-RET; ab134100; 0.1µg/ml; Abcam, Waltham, MA), BDNF (anti-rabbit-BDNF; sc-546; 1:500; Santa Cruz, Dallas, TX) and trkB (anti-goat-TrkB; AF397; 0.5µg/ml; R&D Systems, Minneapolis, MN). Tyrosine hydroxylase protein was quantified using calibrated standard traceable over multiple studies (Salvatore et al., 2022, 2012, 2004; Salvatore and Pruett, 2012). The membranes were then exposed to their respective secondary antibodies and imaged using BioRad imager V3 Chemi-Doc Touch.

The Image Lab software (Bio-Rad Life Sciences, CA) was used for analysis of protein expression. The quantity of proteins of interest (arbitrary values) were expressed as per µg of total protein loaded and normalized based on the ponceau stain intensity in each sample (Eaton et al., 2013; Fosang and Colbran, 2015; Kasanga et al., 2019).

### Immunohistochemistry

TH staining was conducted as previously reported (Kasanga et al., 2022). Briefly, free-floating sections were washed for one day with phosphate-buffered saline (PBS) at 4°C, then permeabilized with PBS + 1% Triton X-100 for 15 min at room temperature. After quenching of endogenous peroxidase with 2.5% H2O2 in 75% methanol, the sections were incubated for 1 h with a blocking solution (4% goat serum + 5% bovine serum albumin + 0.5% Triton X-100). The sections were incubated for 48 h at 4°C with primary mouse anti-TH antibody (anti-mouse-TH; MAB-318, 1:1000; Millipore, Temecula, CA). The sections were then incubated for 1 h with the appropriate secondary antibody after which sections were washed with PBS and incubated with the required detection kits for development. The sections were mounted on glass slides and dehydrated in a series of ethanol solution from 30-100% prior to cover slipping.

For GFR-α1 immunofluorescence staining, 3-4 free-floating sections per hemisphere per subject were washed for 24 h with phosphate-buffered saline (PBS) at 4°C, then permeabilized with 0.5% Triton X-100 for 30 min at room temperature. The sections were incubated for 1 h with a blocking solution (10% bovine serum albumin (BSA), and 0.1% Triton X-100) after washing with PBS. To see which cell types express GFR-α1, GFR-α1 was sequentially stained with TH, and glial fibrillary acidic protein (GFAP). Briefly, antibodies to the three antigens were selected from different species and secondary antibodies labeled with different spectrum of fluorescence were used. Sections were incubated with GFR-α1 antibody (anti-rabbit-GFR-α1; ab216667; 1: 1000; Abcam, Waltham, MA) diluted in 1% BSA at 4°C for 18 h, followed by a 1 h incubation with goat anti-rabbit IgG (H+L) conjugated with Alexa fluor 488 (A32731; 1:1000; Invitrogen, Waltham, MA). The sections were then incubated with GFAP antibody (mouse anti-GFAP; 3670S; 1:1000; CST, Danvers, MA) and TH antibody (sheep anti-TH; 1:1000; AB1542; Millipore Sigma, Burlington, MA) together at 4°C for 18 hours followed by appropriate secondary antibodies (donkey anti-mouse IgG [H+L]; A48257 and donkey anti-sheep IgG [H+L]; A11016; 1:1000; Invitrogen, Waltham, MA) for 1 h. The sections were mounted, and the general background fluorescence was reduced by TrueBlack (TrueBlack® Lipofuscin autofluorescence quencher, Biotium, 23007, CA, US) according to the manufacturer’s instructions.

#### Unbiased stereology

TH+ cells were counted in sections containing the SN pars compacta (SNc) using Stereo Investigator software (MBF Bioscience, USA) and an upright digital microscope (Leica DM4 B Automated Upright Microscope System Cat#DM4B, Leica Biosystems, USA). An optical fractionator was chosen as a stereoscopic probe to measure the size of the cell population. The sampling parameter for both cell counts was set with a counting frame and grid size of 200 × 200 µm and a dissector height of 10 µm.

To quantify TH cells, a contour was created around the SNc region according to The Rat Brain in Stereotaxic Coordinates (Paxinos & Watson, Compact 7th edition, Academic press) using the 4X objective and then counting was performed using the 20X objective. The mounted thickness was measured at each sampling site and the average was calculated to be 11.2µm. Cell counts were estimated from 4-6 sections per sample for TH. The coefficient of error, calculated by the Gundersen formula, was less than 0.1.

#### Immunofluorescence Imaging

Images were acquired using a fluorescence microscope (Keyence, BZ-X800, IL, USA). To analyze the expression of GFR-α1, fluorescence was measured using BZ-X800 analyzer software. Total GFR-α1 expression was calculated by measuring the fluorescence intensity in the SNpc region. For calculating the GFR-α1 expression in TH+ neurons and GFAP+ cells, the target region in the SNpc region was set to either TH+ and GFAP+ cells, and the fluorescence intensity of the GFR-α1 in the target region was measured. The same GFR-α1 fluorescence threshold was used for both measurements. Tissue IHC was normalized by simultaneous staining in one batch, with the same exposure time and threshold applied to all samples according to the fluorescence channel used for imaging fluorescence intensity.

#### Statistical Analysis

The Grubb’s Test was used to detect outliers for each dependent measure with α=0.05 based upon respective *n* obtained for each measure. Results were analyzed using GraphPad Prism 8 (La Jolla, CA, USA) with *p* values < 0.05 considered as significant.

To determine if there was an overall effect of 6-OHDA lesion as compared to the sham-operated group, we first ran a 3-way repeated measures ANOVA (Data reported in Supplemental Table 1), by matching the respective treatment sides (lesioned vs contralateral to lesion in the 6-OHDA group and sham-operated vs intact side in the sham-operated group). Significant differences in treatment side, lesion, or days post-treatment, or any interactions among these variables were reported. To determine differences within the 6-OHDA or sham-operated group, 2-way repeated measures ANOVA was used, matching lesioned (or sham-operated) side to the respective contralateral side results obtained for the 6-OHDA and sham-operation groups, respectively. Time after treatment (sham-operation or lesion), treatment side, or interaction between time and treatment side were reported or the lesion and sham groups respectively. Differences in measures in the same side relative to lesion between the groups segregated by time post-treatment were further evaluated using an unpaired t-test. A paired t-test was used to evaluate differences between hemispheres within a treatment group (7 or 28 days) post-lesion induction or sham-operation; comparing lesioned vs. contralateral hemisphere.

A Pearson correlation was used to evaluate if there was a significant relationship between GFR-α1 expression and TH cell number.

## Results

### Time-dependent loss of nigrostriatal neurons

The 6-OHDA lesion produced a time-dependent loss of TH+ neurons in the SN (Figure 1) (Kasanga et al., 2022). With >90% loss of TH protein confirmed in the striatum (93% by day 7 and 98% by day 28) after lesion induction (Fig. 1A), loss of TH+ neurons in SN progressed less rapidly, with ∼25% loss at day 7 to 80% loss by day 28 (Fig. 1B and C). The results reflect the retrograde and more rapid nature of DA marker loss in striatum compared with SN seen in other preclinical models of PD and human PD (Kneynsberg et al., 2016; Kordower et al., 2013).

**Figure 1.**
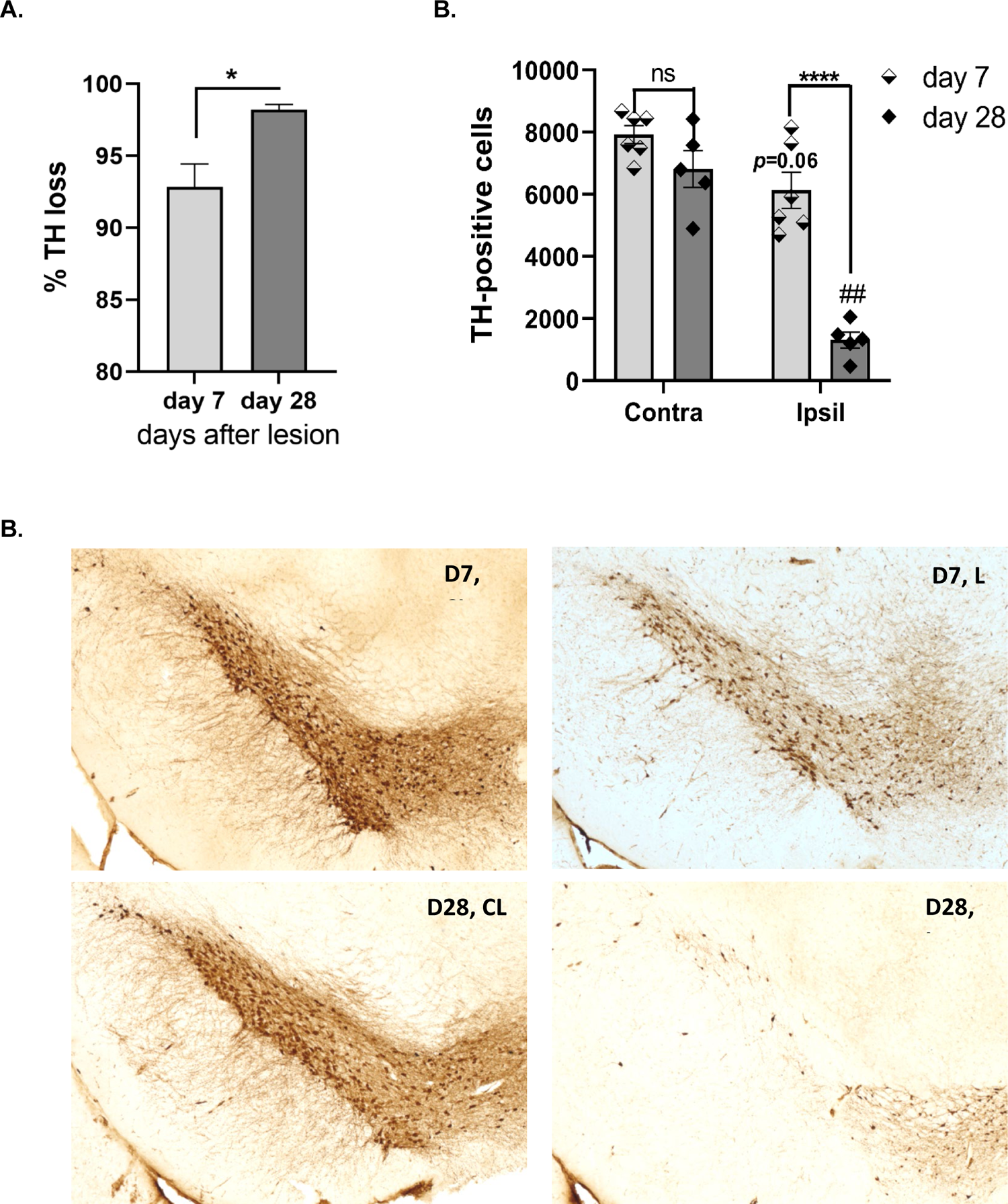
Progressive loss of nigrostriatal neurons following lesion induction by 6-OHDA. **A.** Striatal TH protein loss. TH protein loss exceeded 90% by 7 days after lesion with only slightly greater (4%) loss overall by day 28 (t=2.98, \**p=*0.0155, df=9); **B.** In the same subjects, 6-OHDA lesion produced progressive TH+ neuron loss between 7 and 28 days after induction. Lesion (F_(1,9)_ = 51.4, *p*<0.0001); days post-lesion (F_(1,9)_ = 50.8, *p*<0.0001); lesion x days post-lesion (F_(1,9)_ = 13.3, *p=*0.005). Day 7 vs day 28; lesioned side (t=7.02, \*\*\*\**p*<0.0001, df=9); contralateral to lesioned side (t=1.77, ns, df=9). Contra vs Ipsil to lesion; Day 7 (t=2.43, *p=*0.06, df=5); Day 28 (t=8.16 ^##^*p=*0.001,df=4). **C. Representative image of TH cell loss over time.** Left panels, contralateral to lesion (CL); right panels; ipsilateral to lesion (L). Top panels, Day 7; bottom panels, Day 28. (Kasanga et al., 2022)

### GDNF expression following nigrostriatal lesion

Three-way ANOVA results for GDNF expression in the striatum revealed a significant effect of time (F_(1,28)_=34.96, *p*<0.0001), but not lesion or hemisphere (Supplemental Table 1). There was a significant (25%) lesion-dependent decrease in GDNF expression in the ipsilateral hemisphere 28 days after lesion compared to the contralateral striatum (Fig. 2A). A similar decrease was seen 7 days after lesion. Interestingly, we also observed an increase in GDNF expression in both the ipsilateral and contralateral hemispheres of the 28-day sham cohort (Fig. 2B). As the side contralateral to lesion received a sham-operation in the 6-OHDA group, we speculate the apparent lesion-related decrease may have been due in part to the increase from sham-operation.

**Figure 2.**
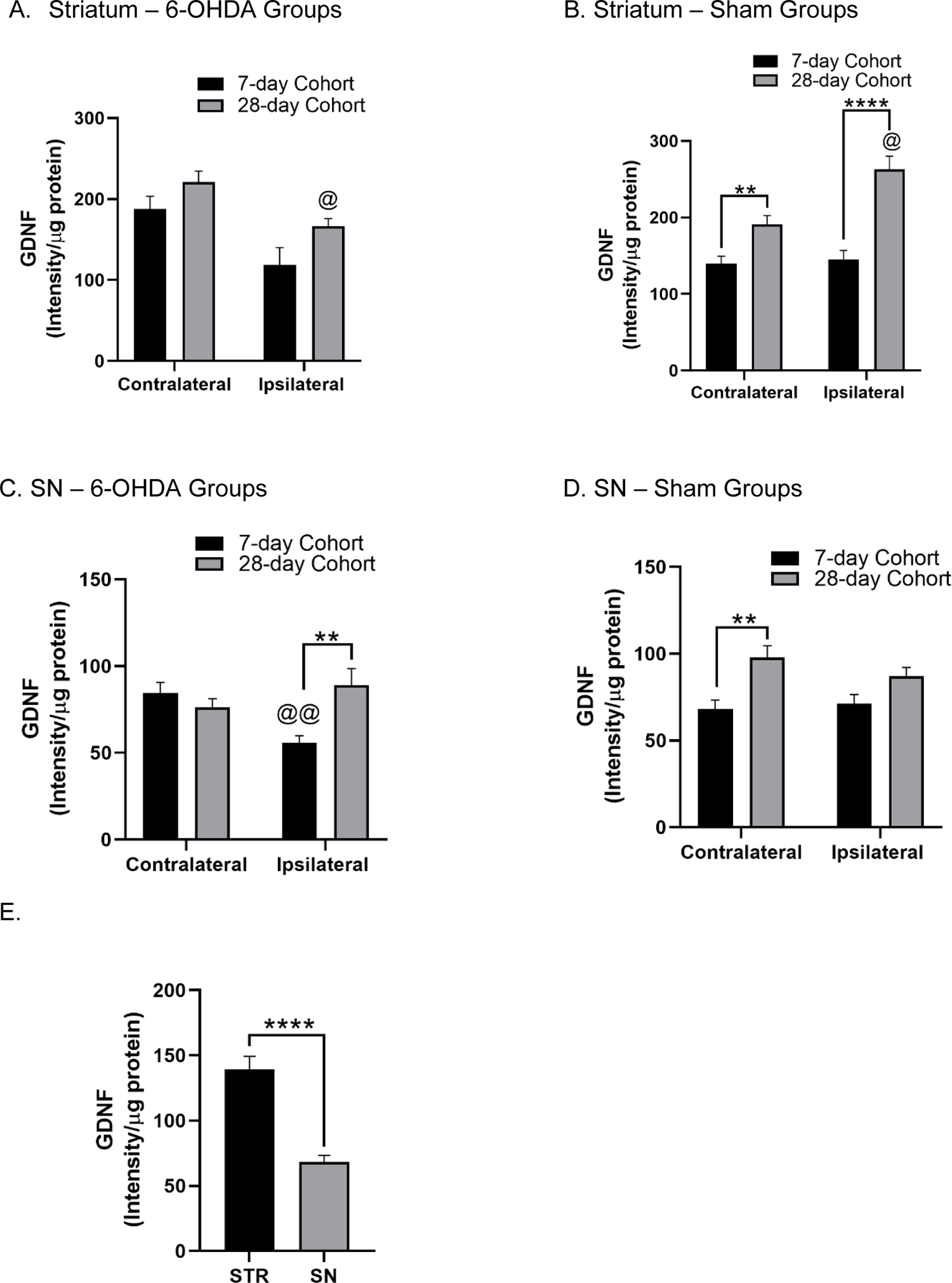
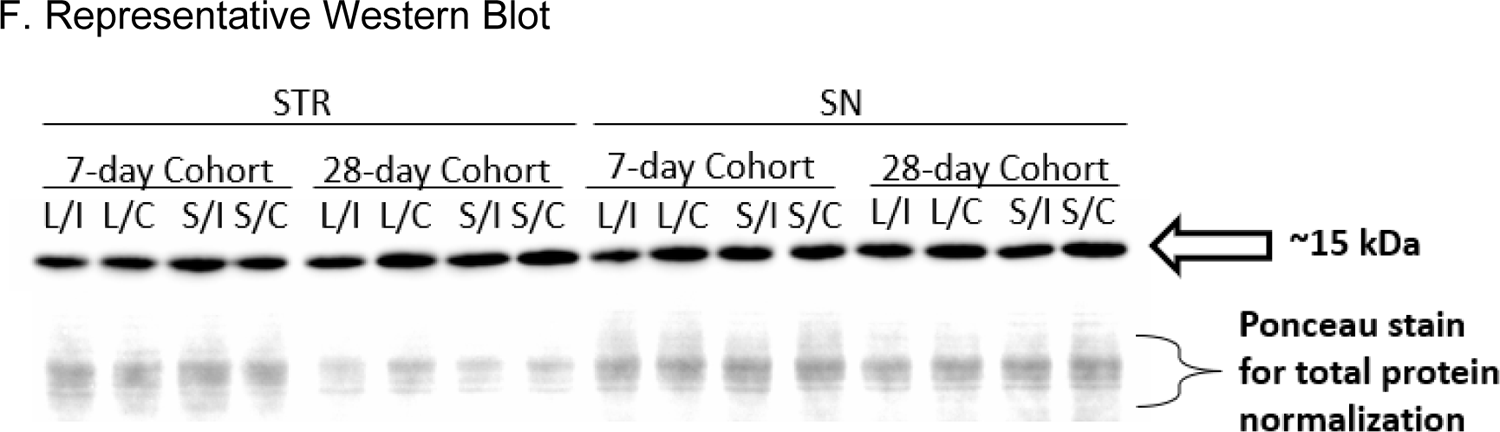
GDNF protein expression in 6-OHDA and sham groups. **(A)** Striatum – 6-OHDA Groups. There was a significant effect of lesion (F_(1, 26)_ = 15.82, *p*<0.001) and time (F_(1, 26)_ = 6.74, *p*<0.05) but there was no lesion x time interaction (F_(1, 26)_ = 0.22, ns). Post-hoc analysis of the main effect revealed a significant decrease in GDNF protein expression in the lesion vs contralateral hemisphere of the 28-day cohort (t=2.868, df=5, @ *p*<0.05). There was a near significant decrease observed in the ipsilateral hemisphere (vs contralateral) of the 7-day cohort (t=2.404, df=6, *p*=0.053). **(B) Striatum – Sham Groups.** Injection of vehicle led to a significant effect of injection (F_(1, 10)_ = 15.68, *p*<0.01), time F_(1, 13)_ = 31.97, *p*<0.0001) and injection x time interaction (F_(1, 10)_ = 12.40, *p*<0.01). Post-hoc analysis revealed a significant increase in the ipsilateral vs. contralateral hemisphere of the 28-day cohort (t=3.230, df=4, @ *p*<0.05). There was also a significant increase in GDNF protein expression in both the ipsilateral (t=5.733, df=12, **** *p*<0.0001) and contralateral (t=3.337, df=11, ** *p*<0.01) hemispheres of the 28-day vs 7-day cohort. **(C) Substantia nigra – 6-OHDA Groups.** There was no significant effect of lesion (F_(1, 29)_ = 1.32, ns) or time F_(1, 29)_ = 3.24, ns) but a significant lesion x time interaction (F_(1, 29)_ = 8.82, *p*<0.01) was observed. Post-hoc analysis revealed a significant decrease in expression in the lesioned vs contralateral hemisphere of the 7-day cohort (t=3.772, df=7, @@ *p*<0.01). A significant decrease in GDNF protein expression was observed in the lesioned hemisphere of the 7-day vs 28-day cohort (t=3.037, df=15, ** *p*<0.01). **(D) Substantia nigra – Sham Groups.** There was a significant effect of time (F_(1, 12)_ = 7.33, *p*<0.05) but not vehicle injection (F_(1, 8)_ = 4.23, ns) or injection x time interaction (F_(1, 8)_ = 3.14, ns). Post-hoc analysis revealed a significant increase in the contralateral hemisphere of the 28-day vs 7-day cohort (t=3.456, df=10, ** *p*<0.01) and a near significant increase in the ipsilateral hemisphere (t=2.068, df=10, *p*=0.0655). **(E) Relative abundance of GDNF.** Striatal and nigral tissues from the 7-day cohort sham group showed significantly greater GDNF expression (∼51%) in the striatum vs SN (t=6.119, df=11, *p*<0.0001). **(F) Representative western blots of GDNF protein expression.** (L- lesion, S- sham, I- ipsilateral, C- contralateral) in the striatum and SN.

Three-way ANOVA results for GDNF expression in the SN depicted a significant effect of time (F_(1,49)_=13.50, *p*=0.0006), but not lesion or hemisphere (Supplemental Table 1). A time-dependent, dichotomous change in GDNF expression was observed. A decrease was seen in the ipsilateral SN following 7-days post-lesioning. This effect was short-lived as levels were restored by day 28 post-lesion (Fig. 2C). There was a significant difference in GDNF expression between the contralateral hemispheres of the 28-day and 7-day cohorts in the sham-operation groups (Fig. 2D). In comparing GDNF protein expression between striatum and SN, we observed a ∼2-fold greater expression level in the striatum (Fig. 2E).

### GFR-α1 expression following nigrostriatal lesion

Three-way ANOVA results for GFR-α1 expression in the striatum revealed a significant effect of lesion (F_(1,29)_=7.36, *p*=0.0111) and hemisphere (F_(1,29)_=28.49, *p*<0.0001; Supplemental Table 1). Nigrostriatal lesion led to a significant decrease in GFR-α1 expression in the ipsilateral striatum (Fig. 2A) with no effect of sham-operation in either the striatum or SN (Fig. 3B, 3D). The lesion impact was observed at both time points (as compared to their respective contralateral hemispheres) with a more significant reduction at day 28 vs day 7 (Fig. 3A), indicating a time-dependent decrease in striatal GFR-α1 expression following lesioning of the nigrostriatal pathway.

**Figure 3.**
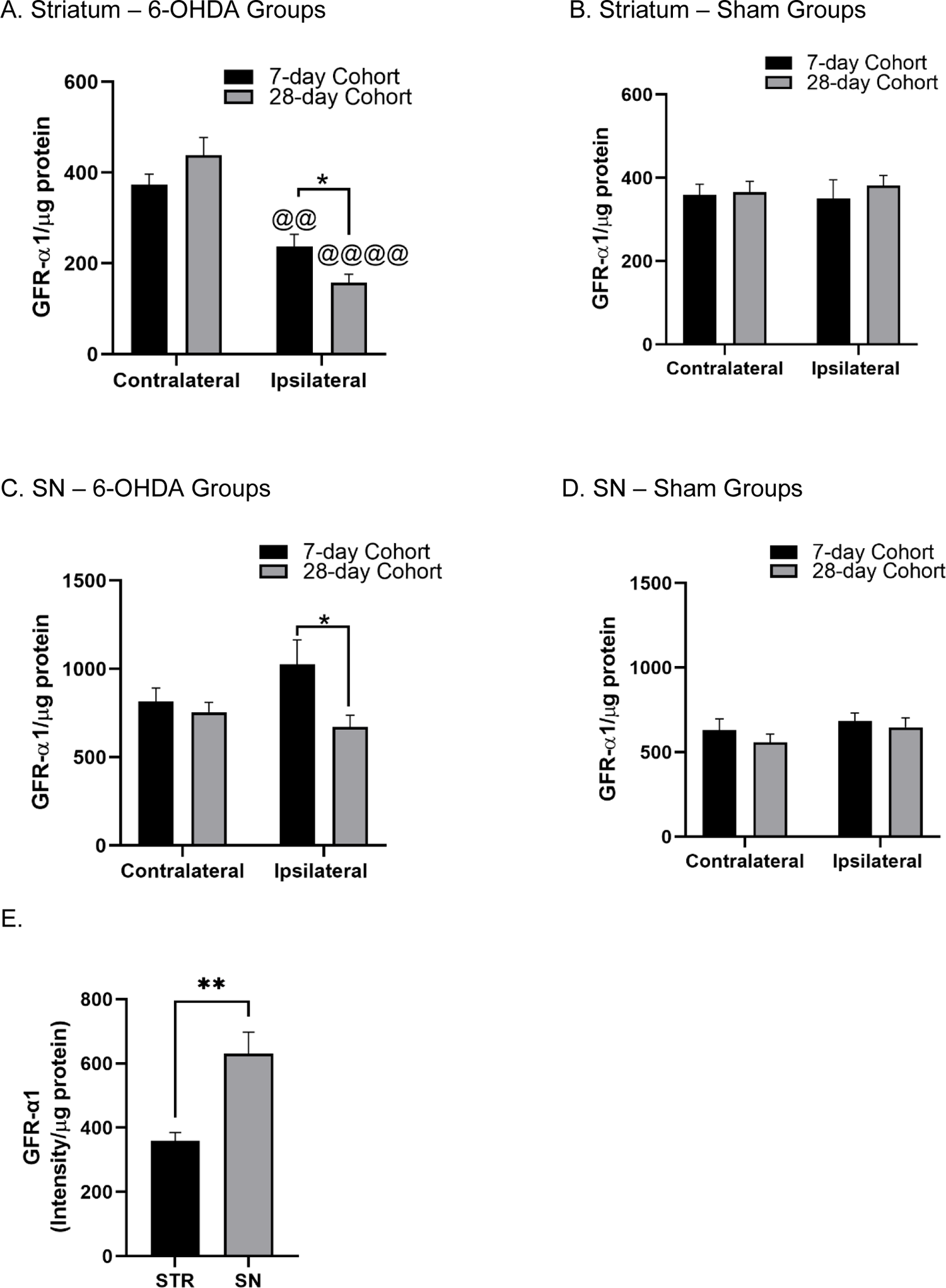
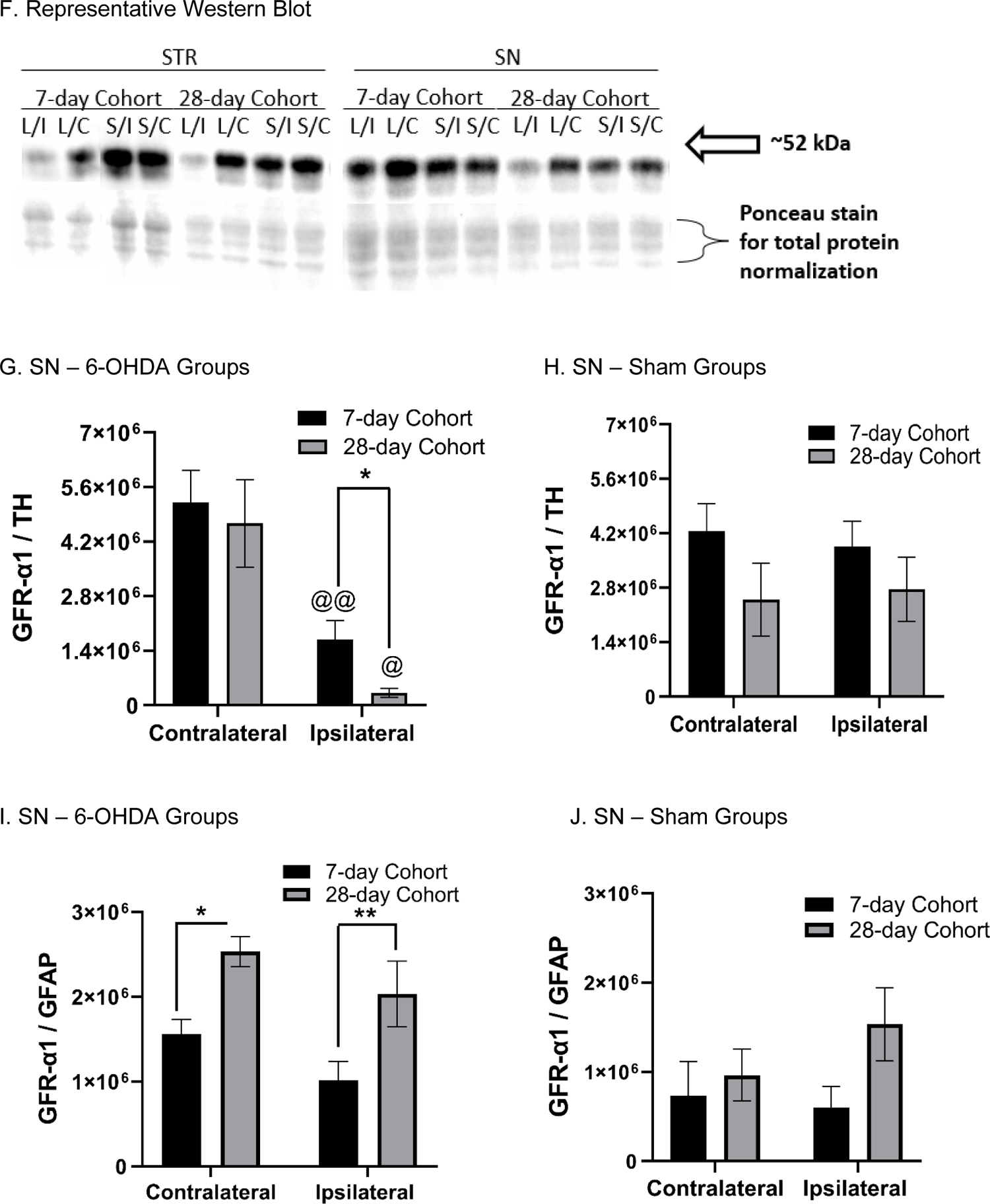
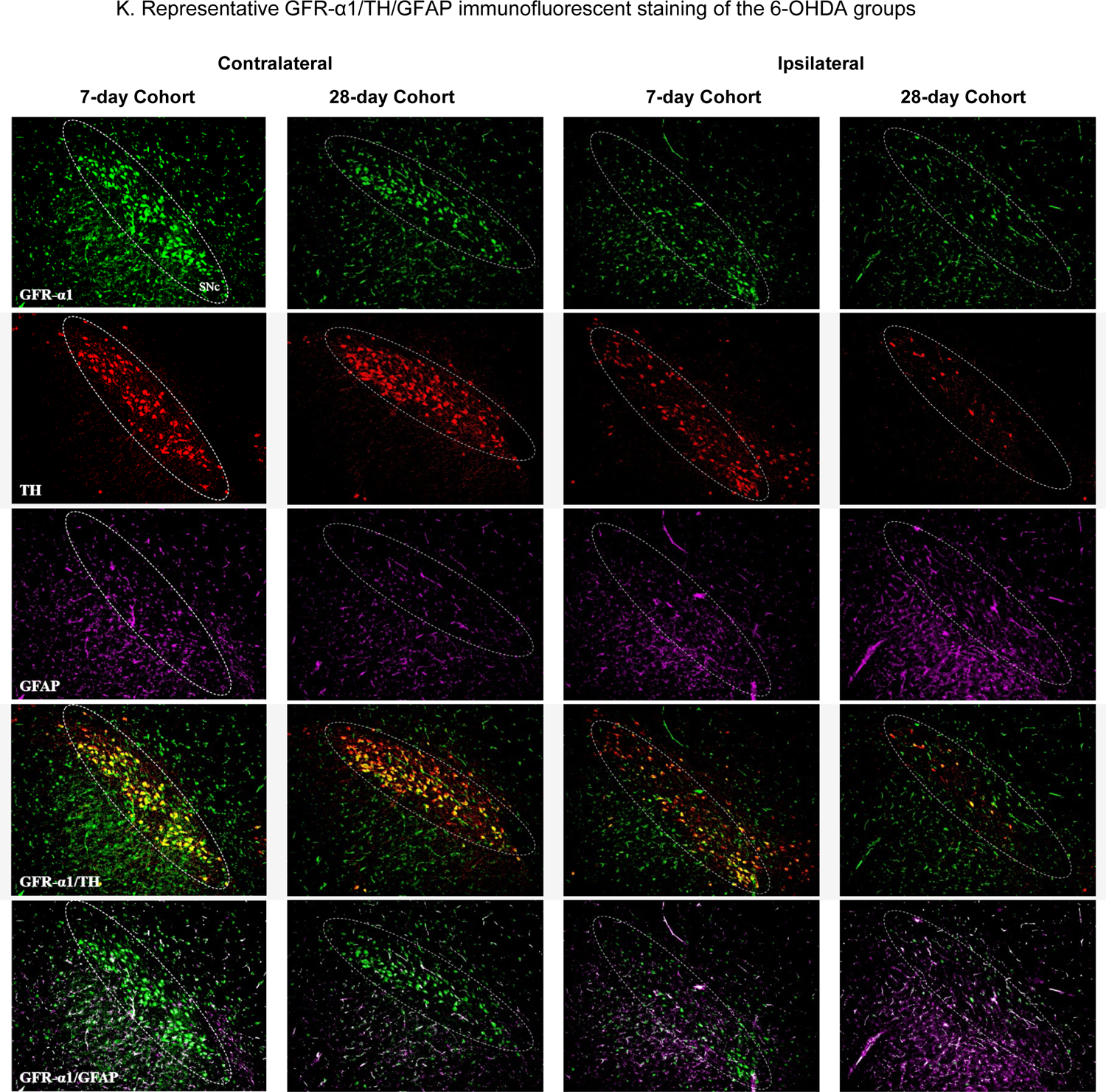
GFR-α1 protein expression in 6-OHDA and sham groups. **(A)** Striatum – 6-OHDA Groups. There was a significant effect of lesion (F_(1, 16)_ = 63.01, *p*<0.0001) and a lesion x time interaction (F_(1, 16)_ = 7.58, *p*<0.05) but there was no significant effect of time (F_(1, 16)_ = 0.06, ns). Post-hoc analysis of the main effect revealed severe loss of GFR-α1 expression in the ipsilateral vs. contralateral hemispheres regardless of duration of lesion (7-day Cohort - t=3.831, df=8, @@ *p*<0.01) and 28-day Cohort - t=7.260, df=8, @@@@ *p*<0.0001). A more significant reduction was observed at day 28 vs day 7 (t=2.431, df=16, * *p*<0.05). **(B) Striatum – Sham Groups.** There was no significant effect of vehicle injection (F_(1, 13)_ = 0.01, ns), time F_(1, 13)_ = 0.26, ns) or injection x time interaction (F_(1, 13)_ = 0.19, ns). **(C) Substantia nigra – 6-OHDA Groups.** There was no significant effect of lesion (F_(1, 29)_ = 0.45, ns) or lesion x time interaction (F_(1, 29)_ = 2.40, ns) but rather a significant effect of time (F_(1, 29)_ = 4.77, *p*<0.05). Post-hoc analysis revealed a significant difference in expression in the lesioned hemispheres of the 7-day vs 28-day cohorts (t=2.303, df=16, * *p*<0.05). **(D) Substantia nigra – Sham Groups.** There was no significant effect of vehicle injection (F_(1, 23)_ = 1.59, ns), time F_(1, 23)_ = 0.94, ns) or injection x time interaction (F_(1, 23)_ = 0.09, ns). **(E) Relative abundance of GFR-α1.** Striatal and nigral tissues from the 7-day cohort sham group showed significantly greater GFR-α1 expression (∼75 %) in the SN vs striatum (t=3.961, df=13, *p*<0.01). **(F) Representative western blots of GFR-α1 protein expression** (L-lesion, S-sham, I-ipsilateral, C-contralateral) in the striatum and SN. **(G) GFR-α1 in TH^+^ neurons – 6-OHDA Groups**. There was a significant effect of lesion F_(1, 9)_ = 44.4, *p*<0.0001) but not time (F_(1, 9)_ = 1.24, ns) or lesion x time interaction (F_(1, 9)_ = 0.50, ns). Post-hoc analysis revealed a significant decrease in GFR-α1/TH co-staining in the lesioned hemispheres of both the 7-(t=5.979, df=5, ** *p*<0.01) and 28-day cohorts (t=3.989, df=4, * *p*<0.05). There was a significant decrease in expression on the lesioned (ipsilateral) **(H) GFR-α1 in TH^+^ neurons – Sham Groups**. There was no significant effect of vehicle injection (F_(1, 8)_ = 0.006, ns), time F_(1, 9)_ = 3.33, ns) or vehicle injection x time interaction (F_(1, 8)_ = 0.17, ns). **(I) GFR-α1 in GFAP^+^ astrocytes – 6-OHDA Groups**. There was a near significant effect of lesion F_(1,15)_ = 4.40, *p*=0.053), and significant effect of time (F_(1, 15)_ = 16.07, *p*=0.001), with no interaction of lesion x time (F_(1, 15)_ = 0.001, ns). Post-hoc analysis revealed a significant increase in GFR-α1/GFAP co-staining in the lesioned hemispheres (t=2.46, \**p*=0.039, df=8) and contralateral to lesion hemispheres (t=3.86, \*\**p*=0.006, df=7) between day 7- and day 28-day cohorts. **(J) GFR-α1 in GFAP^+^ astrocytes – Sham Groups**. There was no significant effect of vehicle injection (F_(1, 7)_ = 0.85, ns), time F_(1,7)_ = 2.01, ns) or vehicle injection x time interaction (F_(1, 7)_ = 2.18, ns). **(K) Representative images of GFR-α1 triple staining.** GFR-α1 expression with TH and GFAP from the contralateral and ipsilateral SN of the 6-OHDA groups.

Three-way ANOVA results for GFR-α1 expression in the SN revealed a significant effect of lesion (F_(1,50)_=12.70, *p*=0.008) and time (F_(1,50)_=6.82, *p*=0.012), with a bilateral increase observed early after lesion in the lesioned groups as compared to the sham groups (Supplemental Table 1). There was a transient increase in GFR-α1 expression in the ipsilateral hemisphere 7 days after nigrostriatal lesion (Fig. 3C).

GFR-α1 protein expression was greater in the SN as compared to the striatum (Fig. 3E). As GFR-α1 is expressed in DA neurons and astrocytes, we evaluated if there was differential expression of GFR-α1 in these cells by IHC. Three-way ANOVA results for GFR-α1/TH revealed a significant effect of time (F_(1,18)_=4.47, *p*<0.05), treatment side (F_(1,17)_=15.5, *p*<0.01), and interaction between lesion and treatment side (F_(1,17)_=14.7, *p*<0.01) (Supplemental Table 1). GFR-α1 expression progressively decreased in TH neurons in the SN ipsilateral to lesion in the day 7- and 28-day cohorts (Fig. 3G). Sham-operation had no effect on GFR-α1/TH co-expression in the SN (Fig. 3H, Supplemental Fig. 1). Three-way ANOVA results for GFR-α1/GFAP revealed a significant effect of lesion (F_(1,15)_=12.7, *p*<0.01), time (F_(1,15)_=11.9, *p*<0.01) (Supplemental Table 1). GFR-α1 expression in GFAP+ cells increased bilaterally as time post-lesion increased (Fig. 3I). Sham-operation had no effect on GFR-α1/GFAP co-expression in the SN, although a trend toward increased expression in the sham-operated side was noted in the 28 day cohort (Fig. 3J, Supplemental Fig. 1).

GFAP immunoreactivity also increased bilaterally in the SN as time past lesion increased. Three-way ANOVA results revealed a significant effect of lesion (F_(1,15)_=9.8, *p*<0.01) and time (F_(1,15)_=9.4, *p*<0.01) (Supplemental Table 1). Sham-operation had no effect on GFAP expression in the SN (data not shown).

There was a significant positive correlation between remaining TH cells and GFR-α1/TH count in the SN (Fig. 4)., suggesting that GFR-α1 expression in TH neurons plays a critical role in nigrostriatal neuron viability. Moreover, the results indicate astrocytes may respond to neuron loss by increasing GFR-α1 expression in response to loss of nigral neurons.

**Figure 4.**
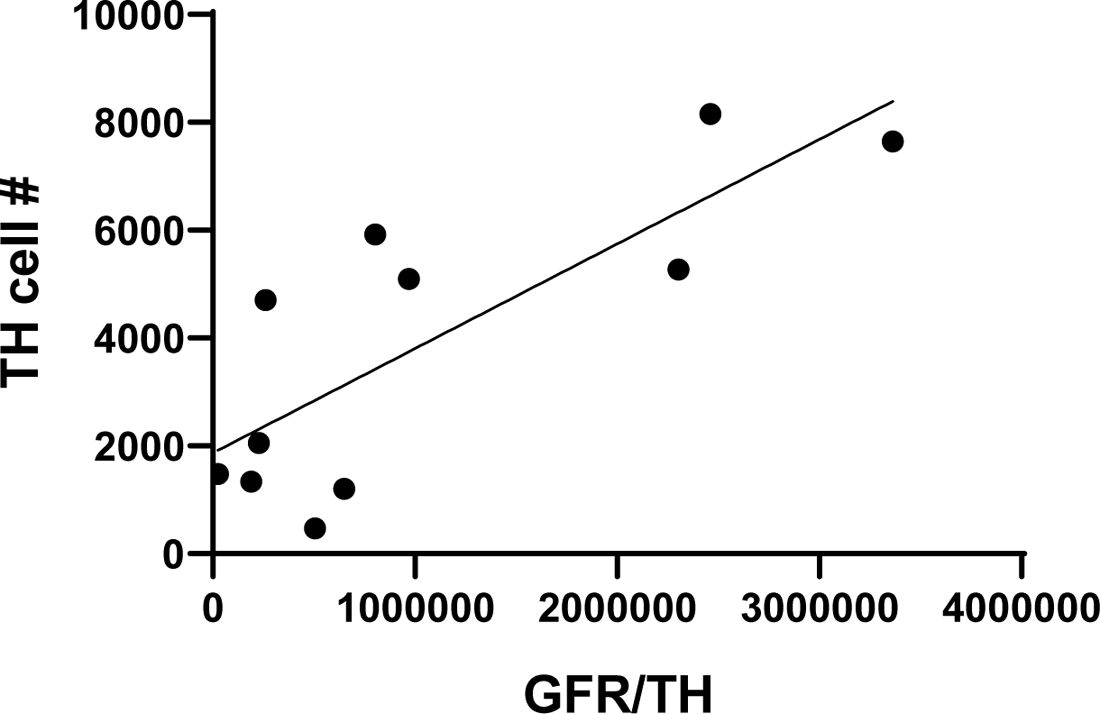
Correlation between GFR-α1 expression in TH^+^ neurons and TH^+^ cell count in subjects with confirmed nigrostriatal lesioning. A positive correlation was found between remaining TH cells and GFR-α1 in TH^+^ neurons count (r=0.7926, ** *p*<0.01).

### RET expression following nigrostriatal lesion

Three-way ANOVA results for RET expression in the striatum revealed a significant effect of lesion (F_(1,27)_=53.03, *p*<0.0001) and hemisphere [F_(1,20)_=86.80, *p*<0.0001; Supplemental Table 1). RET expression in striatum ipsilateral to lesion, was substantially decreased by 7 days. No effect on RET expression occurred in the sham-operation group. (Fig. 5A, 5B). Moreover, lesion also decreased RET expression in the striatum contralateral to lesion by 4 weeks (Fig. 5A).

**Figure 5.**
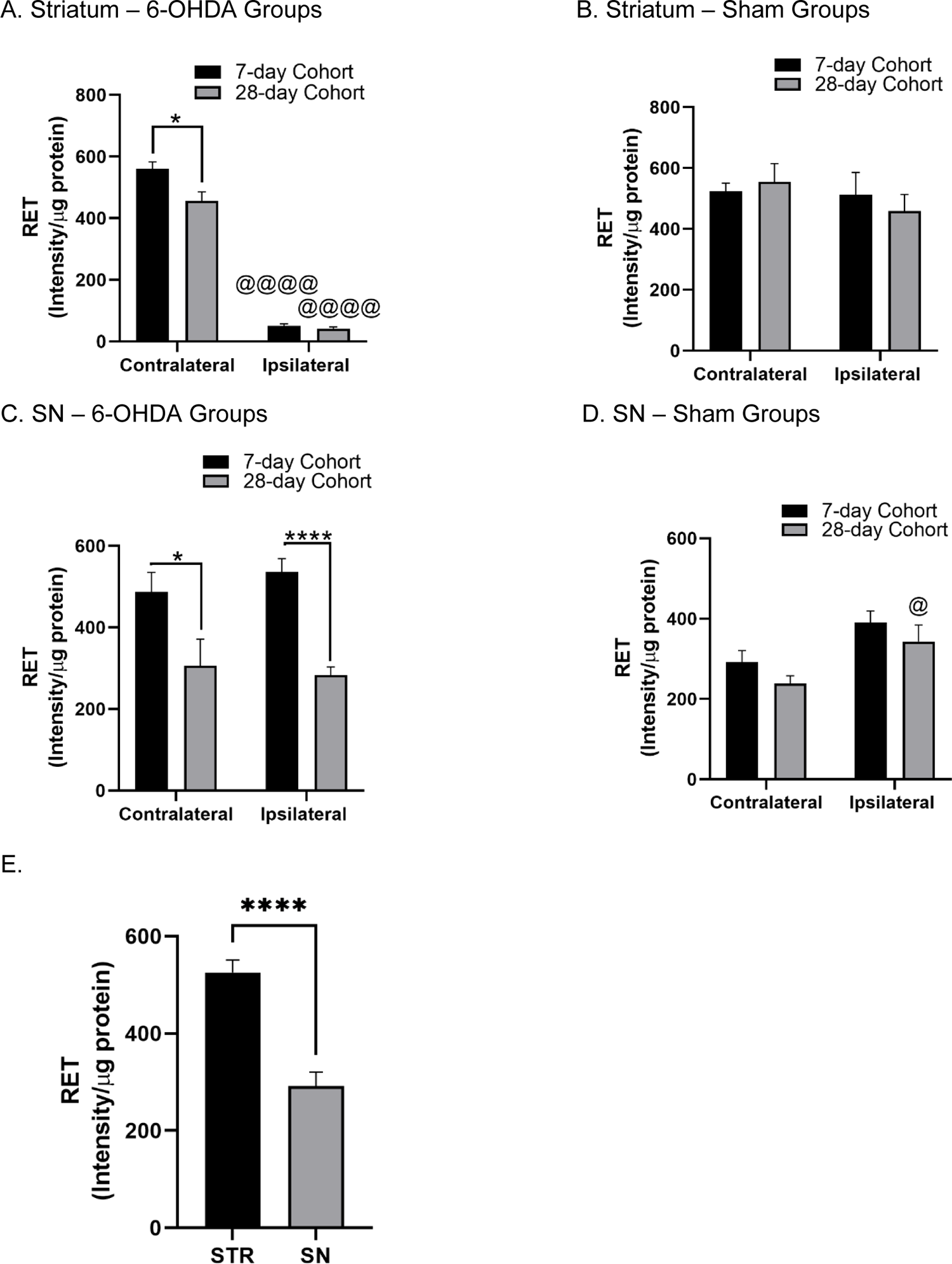
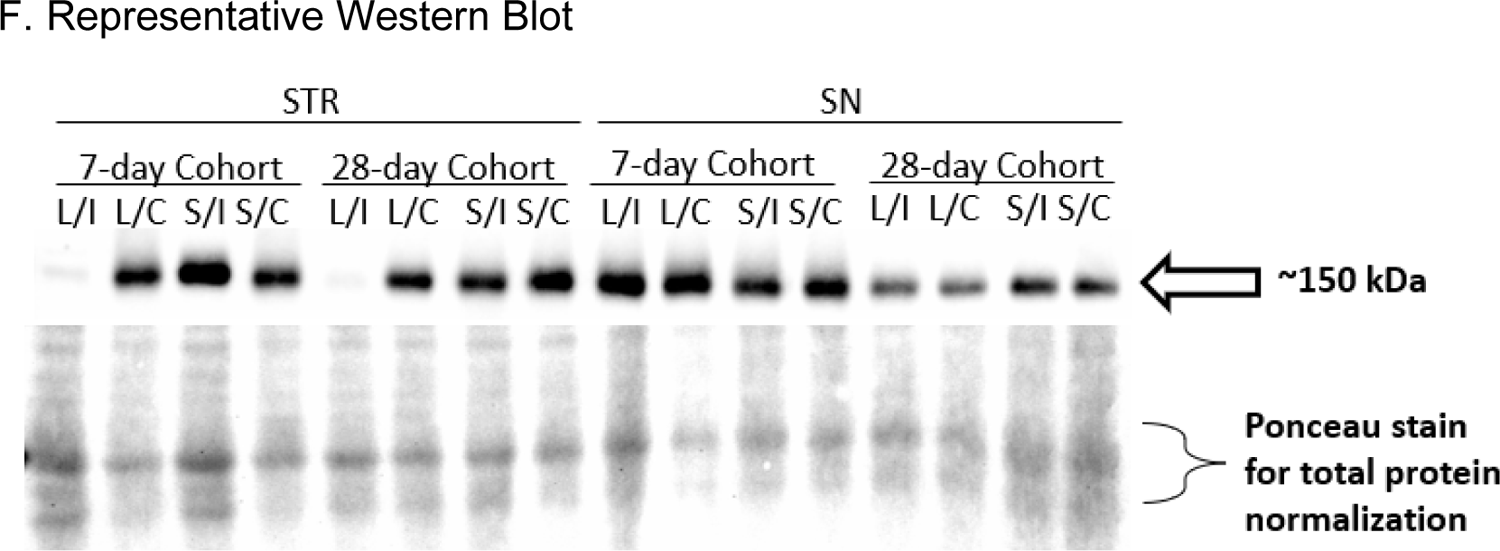
RET protein expression in 6-OHDA and sham groups. **(A)** Striatum – 6-OHDA Groups. Injection of 6-OHDA led to a significant effect of lesion (F_(1, 10)_ = 739.7, *p*<0.0001), time F_(1, 15)_ = 6.97, *p*<0.05) and lesion x time interaction (F_(1, 10)_ = 7.51, *p*<0.05). Post-hoc analysis of the main effect revealed severe loss of RET expression in the lesioned vs. contralateral hemispheres regardless of duration of lesion (7-day Cohort - t=21.26, df=5, @@@@ *p*<0.0001) and 28-day Cohort - t=15.21, df=5, @@@@ *p*<0.0001). There was also a significant decrease in RET protein expression in the contralateral hemisphere of the 28-day vs 7-day cohort (t=2.819, df=13, * *p*<0.05). **(B) Striatum – Sham Groups.** There was no significant effect of vehicle injection (F_(1, 10)_ = 1.01, ns), time F_(1, 12)_ = 0.05, ns) or injection x time interaction (F_(1, 10)_ = 0.50, ns). **(C) Substantia nigra – 6-OHDA Groups.** There was no significant effect of lesion (F_(1, 11)_ = 0.08, ns) or lesion x time interaction (F_(1, 11)_ = 0.83, ns) but rather a significant effect of time (F_(1, 14)_ = 18.85, *p*<0.001). Post-hoc analysis revealed a significant decrease in RET expression in the lesioned (t=6.508, df=13, **** *p*<0.0001) and contralateral (t=2.249, df=12, * *p*<0.05) hemispheres of the 28-day vs 7-day cohorts **(D) Substantia nigra – Sham Groups.** There was a significant effect of vehicle injection (F_(1, 10)_ = 12.440, *p*<0.01) but not time (F_(1, 13)_ = 2.011, ns) or injection x time interaction (F_(1, 10)_ = 0.003, ns). Post-hoc analysis of the main effect revealed a significant increase in RET protein expression in the lesioned hemispheres vs contralateral hemispheres of the 28-day cohort (t=2.606, df=5, @ *p*<0.05) and a near significant increase in the 7-day cohort (t=2.061, df=5, *p*=0.0944). **(E) Relative abundance of RET.** Striatal and nigral tissues from the 7-day cohort sham group showed significantly greater GDNF expression in the striatum vs SN (t=5.950, df=11, *p*<0.0001). **(F) Representative western blots of RET protein expression.** (L- lesion, S- sham, I- ipsilateral, C- contralateral) in the striatum and SN. **(G)** Representative images of RET staining with TH from the contralateral and ipsilateral SN of the 6-OHDA groups (SNc, substantia nigra pars compacta; SNr, substantia nigra pars reticulata).

Three-way ANOVA results for RET expression in the SN revealed a significant effect of lesion (F_(1,28)_=6.82, *p*<0.05) and time (F_(1,28)_=4.67, *p*<0.05), with a bilateral increase observed early after lesion in the lesioned groups as compared to the sham groups (Supplemental Table 1). The bilateral increase in RET at 7 days post-lesion returned to levels in the sham-op group by 28 days (Fig. 5C). This persistence of RET expression in the remaining dopaminergic neurons has been reported in clinical cases of PD (Chu and Kordower, 2021; Walker et al., 1998). There was minimal effect of sham-operation on RET expression, thus indicating lesion-related increases at day 7 were specific processes associated with eventual loss of nigrostriatal neurons (Fig. 5D). Interestingly, RET protein expression was greater in the striatum than in SN (Fig. 5E), which was the converse of differences in relative expression of GFR-α1 between these regions.

### BDNF and TrkB expression following nigrostriatal lesion

Unlike the pronounced effect of nigrostriatal loss seen with GDNF receptors, RET and GFR-α1 (Figs. 2-5), there was no significant effect of nigrostriatal lesioning on total BDNF or TrkB protein expression (Figs. 6-7; Supplemental Table 1). In the sham-op group, comparing striatal vs nigral levels, BDNF protein expression was significantly higher in the SN vs striatum (Fig. 6E). In contrast, TrkB protein expression was higher in the striatum vs. SN (Fig. 7E).

**Figure 6.**
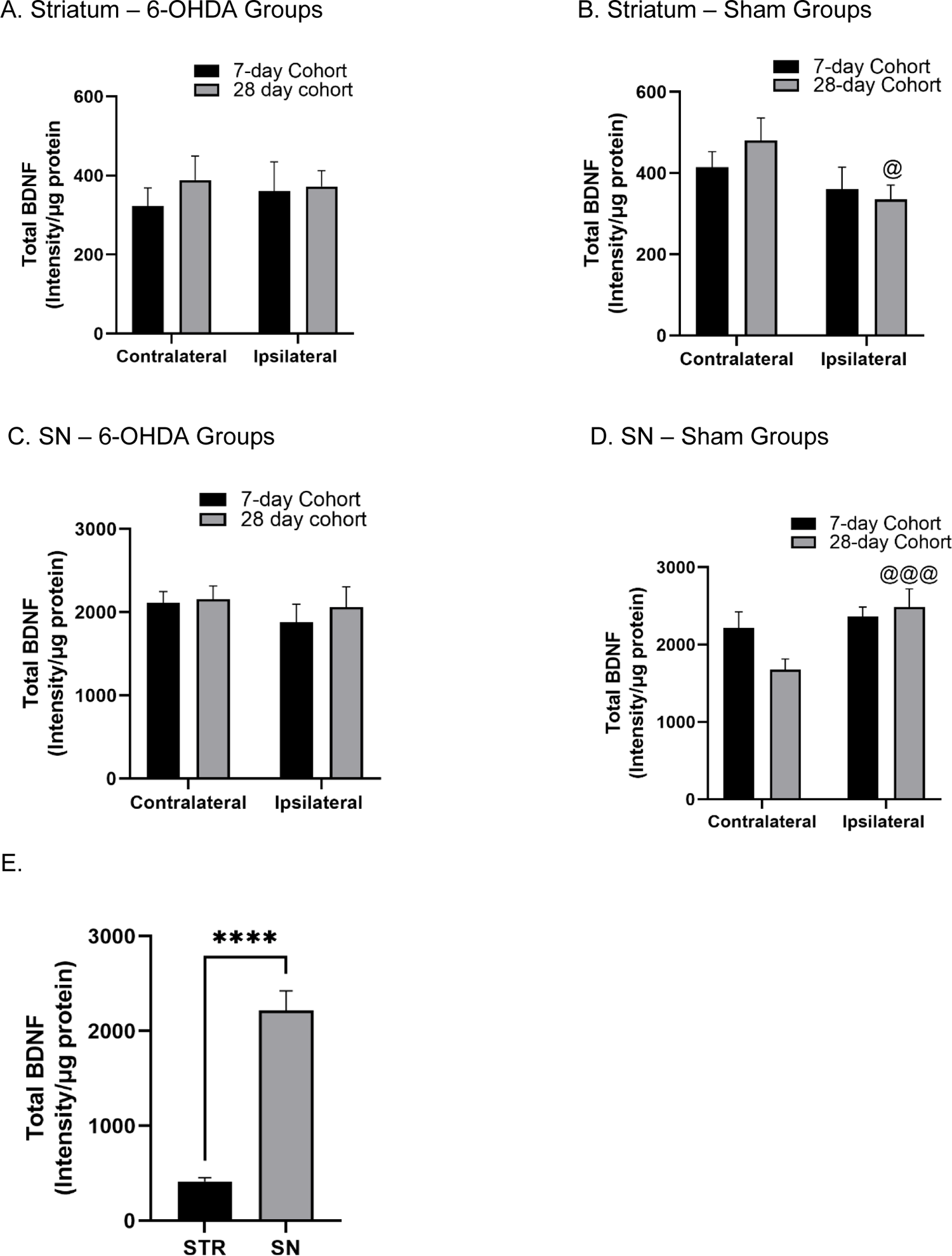
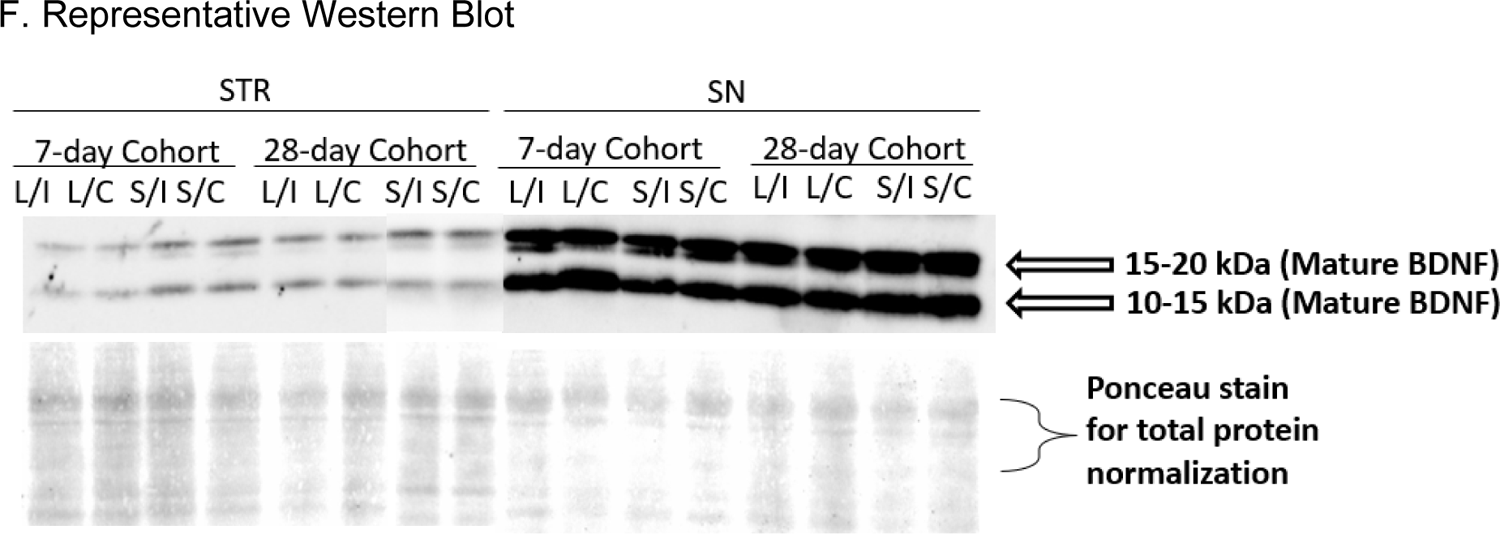
Total BDNF precursor protein expression in 6-OHDA and sham groups. **(A)** Striatum – 6-OHDA Groups. There was no significant effect of lesion (F_(1, 15)_ = 0.080, ns), time (F_(1, 16)_ = 0.326, ns) or lesion x time interaction (F_(1, 15)_ = 0.381, ns). **(B) Striatum – Sham Groups.** There was a significant effect of vehicle injection (F_(1, 10)_= 6.63, *p*<0.05) but not time F_(1, 13)_ = 0.308, ns) or injection x time interaction (F_(1, 10)_ = 2.319, ns). Post-hoc analysis of the main effect revealed a significant decrease in total BDNF protein expression in the ipsilateral hemisphere vs contralateral hemispheres of the 28-day (t=2.918, df=4, @ *p*<0.05). **(C) Substantia nigra – 6-OHDA Groups.** There was no significant effect of lesion (F_(1, 16)_ = 1.172, ns), time (F_(1, 16)_ = 0.254, ns) or lesion x time interaction (F_(1, 16)_ = 0.196, ns). **(D) Substantia nigra – Sham Groups.** There was a significant effect of vehicle injection (F_(1, 13)_ = 12.58, *p*<0.01) and injection x time interaction (F_(1, 13)_ = 6.02, *p*<0.05) but not time F_(1, 13)_ = 0.93, ns). Post-hoc analysis of the main effect revealed a significant decrease in total BDNF protein expression in the contralateral hemisphere (vs ipsilateral hemisphere) of the 28-day cohort (t=5.985, df=6, @@@ *p*=0.001) and a near significant decrease between the contralateral hemispheres of the 7- and 28-day cohorts (t=2.109, df=13, *p*=0.0550). **(E) Relative abundance of BDNF.** Striatal and nigral tissues from the 7-day cohort sham group showed significantly greater BDNF expression in the SN vs striatum (t=8.014, df=13, *p*<0.0001). **(F)** Representative western blots of BDNF protein expression (L- lesion, S- sham, I- ipsilateral, C- contralateral) in the striatum and SN.

**Figure 7.**
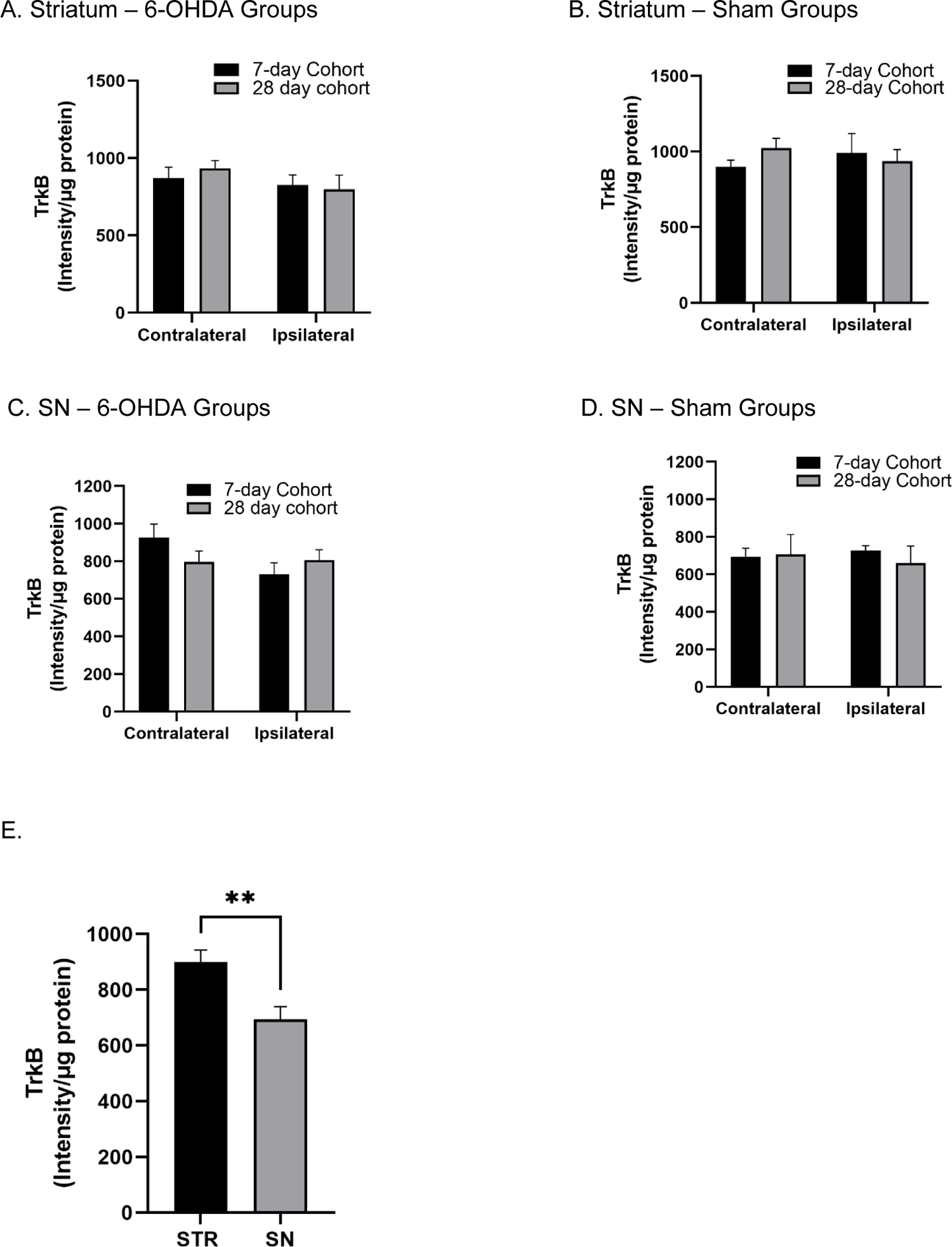
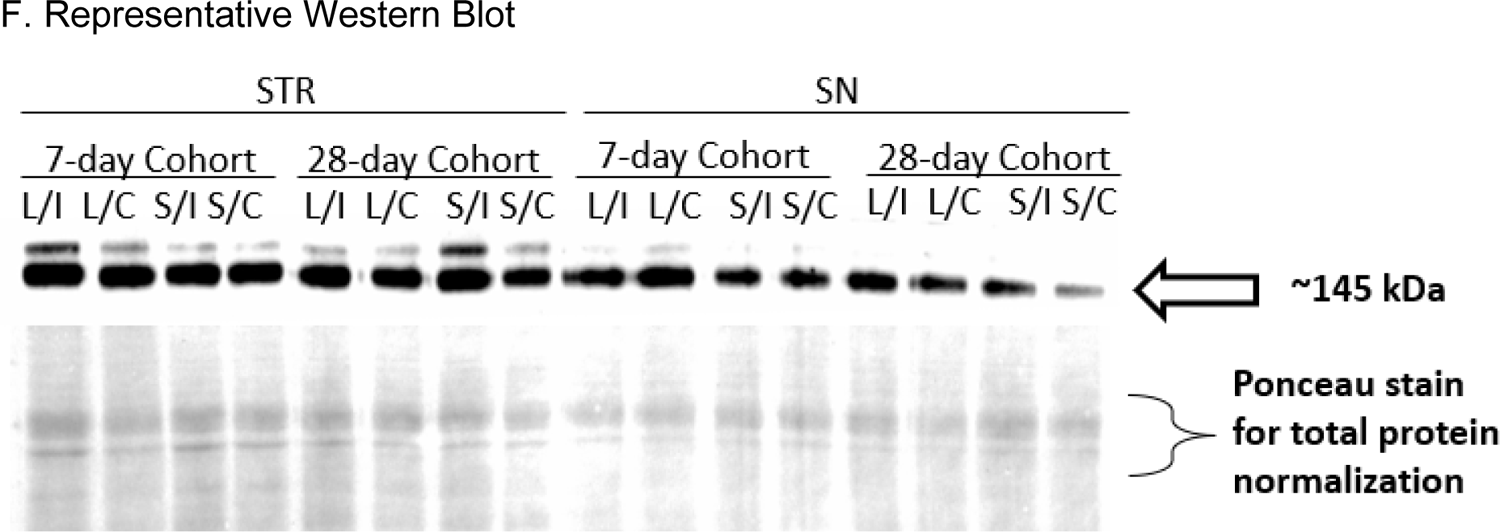
TrkB protein expression in 6-OHDA and sham groups. **(A)** Striatum – 6-OHDA Groups. There was no significant effect of lesion (F_(1, 11)_ = 2.19, ns), time (F_(1, 16)_ = 0.0004, ns) or lesion x time interaction (F_(1, 11)_ = 1.31, ns). **(B) Striatum – Sham Groups.** There was no significant effect of vehicle injection (F_(1, 9)_ = 0.00007, ns), time F_(1, 9)_ = 0.19, ns) or injection x time interaction (F_(1, 9)_ = 0.92, ns). **(C). Substantia nigra – 6-OHDA Groups** There was no significant effect of lesion (F_(1, 13)_ = 2.46, ns), time (F_(1, 15)_ = 0.13, ns) or lesion x time interaction (F_(1, 13)_ = 2.99, ns). **(D) Substantia nigra – Sham Groups.** There was no significant effect of vehicle injection (F_(1, 8)_ = 1.51, ns), time F_(1, 12)_ = 0.04, ns) or injection x time interaction (F_(1, 8)_ = 1.19, ns). **(E)** S **Relative abundance of TrkB.** Striatal and nigral tissues from the 7-day cohort sham group showed significantly greater TrkB expression (∼30 %) in the striatum vs SN (t=3.201, df=11, *p*<0.01). **(F)** Representative western blots of TrkB protein expression (L- lesion, S- sham, I- ipsilateral, C- contralateral) in the striatum and SN.

## Discussion

Preclinical results with GDNF provided significant impetus to initiate clinical trials. Despite evidence of improved motor function in early open-label clinical trials with GDNF (Gill et al., 2003; Slevin et al., 2005), subsequent placebo-controlled trials showed primary endpoints were not met (Lang et al., 2006; Whone et al., 2019). Our results strongly point to the possibility that the lack of available GDNF receptors, particularly GFR-α1, on nigrostriatal neurons may be a significant barrier to GDNF therapeutic efficacy. As the temporal sequence of GDNF or BDNF receptor expression between striatum and SN during nigrostriatal neuron loss has not been established, we determined that expression of the ligands, GDNF and BDNF, were largely unaffected by nigrostriatal neuron loss, and, instead show that the GDNF receptors, GFR-α1 and RET, were differentially affected between striatum and SN over time after lesion induction. Of note, the samples used in this study were obtained from rats that had confirmed parkinsonian motor phenotypes, and significant loss of TH protein in both striatum (>95%) and SN (∼60%) by day 7 and even greater loss (>80%) in SN by day 28 (Kasanga et al., 2022). Notably, GDNF expression was minimally affected after lesion at both time points. In contrast, both RET and GFR-α1 expression levels decreased differentially between the striatum and SN as lesion progressed. However, expression of the BDNF receptor, TrkB, did not change in either striatum or SN at any time point evaluated after lesion induction. The results in our study clearly point to the likelihood that the progressive loss of GFR-α1, and possibly RET in the striatum, are critical barriers to overcome to improve GDNF efficacy as nigrostriatal neuron loss progresses. Importantly, our findings offer a possible explanation for failure to meet primary endpoints despite extended GDNF delivery (Whone et al., 2019), given that clinical trials administered GDNF well past 5 years of PD diagnosis; a time period where DA markers in striatum are largely absent (Kordower et al., 2013).

Behavioral improvement and improved DA signaling in preclinical studies occurred when GDNF was delivered during progression of nigrostriatal lesion, or in aging models (Bowencamp et al., Gash et al., 1996; Gerhardt et al., 1999; Grondin et al., 2003; Salvatore et al., 2004, 2009). It may be argued that the timing of delivery in preclinical studies did not fully represent the severity of the disease stage in the clinical trials, wherein loss nigrostriatal neuron terminals was likely complete when GDNF was initiated ∼8 years post-diagnosis. The rats used in this study had greater than 95% TH and DA loss in striatum and beginning of nigrostriatal cell body loss and motor impairment (Kasanga et al., 2022), which gives a perspective on the possibility that the patients in clinical trials likely had very little, if any, available RET or GFR-α1. Few preclinical studies have investigated neuroprotection or restoration by targeting the GDNF receptors, but the available evidence suggests plausibility from such an approach to augment DA signaling by preserving TH protein expression in the SN (Kasanga et al., 2019; Pruett and Salvatore, 2013). Therefore, this study represents a new launch point to exploit the receptor components of GDNF signaling as targets for attenuating nigrostriatal neuron loss and improve motor function in PD.

Our study did not reveal any significant change in BDNF expression or its cognate receptor, TrkB. This lack of effect was somewhat unexpected, given earlier work indicating that BDNF signaling may be implicated in the pathogenesis of PD (Gall et al., 1992; Hyman et al., 1991). For instance, decreased BDNF expression (both peripherally and centrally) and phosphorylated TrkB levels have been reported in PD patients (Huang et al., 2019; Jiang et al., 2019; Mogi et al., 1999; Parain et al., 1999). Consistent with our findings, however, RET, but not TrkB, ablation causes progressive and adult-onset loss of DA neurons specifically in the SN (Kramer et al., 2007), and targeted disruption of the BDNF gene does not induce dopaminergic cell loss or terminal projections to the striatum (Jones et al., 1994) or affect DA tissue levels (Bastioli et al., 2022).. In contrast, some studies do show that loss of TrkB induces some moderate loss of nigrostriatal neurons (Halbach et al., 2005), or causes a modest increase in MPTP-induced neuron loss (Baydyuk et al., 2011). As such, TrkB agonists may be promising candidates in preclinical PD models (Jin, 2020; Nie et al., 2015; Sconce et al., 2015). Unlike GDNF, BDNF has not been tested in the clinic, and majority of the evidence supporting that BDNF affects DA signaling originate from TrkB agonist studies (Chmielarz and Saarma, 2020; Palasz et al., 2020; Tian et al., 2022). In summary, the relationship of BDNF signaling with nigrostriatal viability and function is still yet unclear. However, our results would suggest that any influence of BDNF signaling on nigrostriatal neuron function may be well before the severe stages of nigrostriatal terminal loss.

Receptor-mediated trafficking of exogenous GDNF and receptor-mediated transduction of GDNF signaling to the soma has been argued to be a critical mechanism for GDNF efficacy in PD treatment (Ito and Enomoto, 2016; Leitner et al., 1999; Tansey et al., 2000; Tomac et al., 1995; Zahavi et al., 2015). The substantial loss of RET and gradual loss GFR-α1 in the striatum, as shown by our results, would portend minimal engagement of GDNF signaling for downstream signal transduction (Barroso-Chinea et al., 2005), even if GDNF is given at the time of PD diagnosis. This is also supported by clinical data where diminished RET expression in nigral neurons evaluated in post-mortem SN had undetectable levels of TH and phosphor-ribosomal protein S6, a useful indicator of pathway activation and neuronal activity (Biever et al., 2015; Chu and Kordower, 2021). Notably, these observations were from subjects that reported minimal motor deficits, suggesting that impaired RET signaling may begin in patients even in the prodromal stages of PD (Chu and Kordower, 2021). On the other hand, our results also point to possible plasticity in GDNF signaling in the SN, and RET may be a viable target to amplify GDNF signaling. Despite overall nigrostriatal neuron loss 7 days after lesion induction (wherein striatal TH loss already exceeded 90%), RET expression at that time showed a transient bilateral increase that was isolated within the SN; loss in striatum was already maximal at the same time point. Based upon earlier work, this nigra-specific increase in RET expression may represent an inherent CNS response to maintain DA signaling against unrelenting nigrostriatal neuron loss (Drinkut et al., 2016). Clearly, more investigation in the relative influence of RET expression in striatum and SN may finally resolve where in the nigrostriatal pathway it may have the greatest influence on DA signaling and neuron viability. It also suggests, given that GDNF levels are minimally affected during nigrostriatal neuron loss, that augmenting RET itself may prove more efficacious to augment GDNF signaling.

In the CNS, GFR-α1 exists as glycosylphosphatidylinositol-linked insoluble and secreted soluble forms (Paratcha et al., 2001; Pruett and Salvatore, 2010), and it has been detected in both neuronal and non-neuronal cells (Luis and Pascual, 2016; Marco et al., 2002; Pruett and Salvatore, 2013; Remy et al., 2001; Sarabi et al., 2001). In fact, the soluble form of GFR-α1 is expressed and secreted by glial cells (Paratcha et al., 2001). Our immunohistochemical analysis revealed progressive decreases in GFR-α1/TH co-expression in the ipsilateral SN of the lesioned groups (Fig. 3G) along with a bilateral increase in GFR-α1/GFAP co-expression with lesion progression (Fig. 3I). GFR-α1 is predominantly localized in neuronal cells in the intact rodent nigrostriatal system (Luis and Pascual, 2016; Marco et al., 2002). It is also responsive to CNS injury, and in line with our results showing increased expression in astrocytes of lesioned, but not sham-operated, rats, there are several reports of increased GFR-α1 in glial cells in excitotoxic conditions or following mechanical injury (Bresjanac and Antauer, 2000; Marco et al., 2002; Remy et al., 2001; Widenfalk et al., 2001). It has been suggested that dopaminergic neurons are the major source of cells in the midbrain expressing RET (Kramer and Liss, 2015). However, our results show a strong correlation of GFR-α1 with TH neuron number, supporting the possibility that GFR-α1 expression is equally, if not more, critical for trophic support of nigrostriatal neurons. It is feasible to determine if increased expression of GFR-α1 by the astrocytes may be a CNS response to forestall the loss of TH+ neurons, as the loss of GFR-α1 in TH neurons was countered with increased GFR-α1 expression in GFAP+ cells (likely astrocytes). Taken together, GDNF signaling may be greatly impeded by the significant loss in GFR-α1 in both the terminals and cell bodies of nigrostriatal neurons, an effect that may be partly counteracted by an upregulation of GFR-α1 by astrocytes in the SN. Thus, it is feasible that the increased expression of GFR-α1 in the astrocytes, as the soluble form of GFR-α1, may signal in *trans* on adjacent remaining DA neurons to provide trophic support along with remaining RET. However, this possibility has yet to be investigated.

### Summary

For over two decades, the extent of GDNF’s efficacy to alleviate or partially reverse motor impairment in PD clinical trials has been the topic of considerable debate (Barker et al., 2020; Manfredsson et al, 2020). One of the prevailing questions for alleviating motor impairment in PD with GDNF is whether the clinical trials that failed to demonstrate efficacy are true failures or false failures. True failures as depicted by a true negative outcome from the clinical trial versus a false failure which could be a failure due to methodology (Espay et al., 2020). In this case, if the delivery method of GDNF is a reason for failure due to poor distribution in the CNS (Ai et al., 2003; Salvatore et al., 2006), there is the inherent assumption that there are receptors available for GDNF for it to exert trophic effects. Our results strongly suggest that lack of efficacy may not be a question of adequate GDNF levels, but severe loss of the receptors that would bind it; RET and GFR-α1, at least in the striatum. We show the GDNF signaling receptors RET and GFR-α1, begin to decrease substantially early in the striatum, and GFR-α1 loss on nigrostriatal cell bodies in the SN correlates with nigrostriatal neuron loss. This loss, while not causing complete TH protein and neuron loss (Kasanga et al., 2022), was still critically tied to neuron viability. Such loss would arguably impact the efficacy of endogenous or exogenous GDNF. Our finding sheds new light on plausible reasons for the inconsistent results generated in GDNF clinical trials. They also lend credence to the argument that GDNF would be effective as treatment for PD if one of these therapeutic strategies are considered; 1) deliver GDNF early enough to capture sufficient GFR-α1 levels remaining in striatum, particularly in the prodromal period of PD, 2) augment GFR-α1 expression in the SN, wherein along with still detectable levels of RET, endogenous GDNF could be sufficient to drive recovery of TH protein or promote survival of remaining nigral dopaminergic neurons, or 3) augment RET and GFR-α1 expression in the striatum to enable retrograde trafficking of the ligand-receptor complex back to the SN. These possibilities are supported by our study, and provide feasible and logical rationale why GDNF treatment for PD should not be abandoned at this time.

## Supporting information

Supplemental Results

## Authors’ contributions

Study design; MFS, JRR, EAK. Study conduct; EAK, YH, MKS. Data collection and analysis; MFS, JRR, EAK, YH, MKS, CP, AB, WN, RM. Interpretation of results; MFS, JRR, FPM, EAK, YH. Writing manuscript; MFS, EAK. Editing manuscript; MFS, JRR, FPM, VAN, EAK. All authors read and approved the final manuscript.

## Acknowledgements

This work was supported by the Department of Defense Parkinson’s Research Program, Investigator-Initiated Research Award (W81XWH-19-1-0757) awarded to MFS. EAK was supported by the Office of Vice President for Research and Innovation, the Institute for Healthy Aging, and National Institutes of Health/National Institute on Aging Predoctoral International Fellowship (T32 AG020494), and the Parkinson’s Foundation Visiting Scholar Award. Additional support was provided by R01ES033892 and U01NS108956 to JRR and a CV Starr Fellowship to AB. The funders had no role in the study design, collection, analysis, and interpretation of data, writing of the manuscript, or decision to submit the article for publication.

## Data Availability

Data generated or analyzed during this study are included in this published article and its supplementary information files. Additional data are available from the corresponding author on reasonable request.

