## Supplemental Results for "Differential expression of RET and GDNF family receptor, GFR-α1, between striatum and substantia nigra following nigrostriatal lesion: a case for diminished GDNF-signaling"

### Supplementary Figure 1

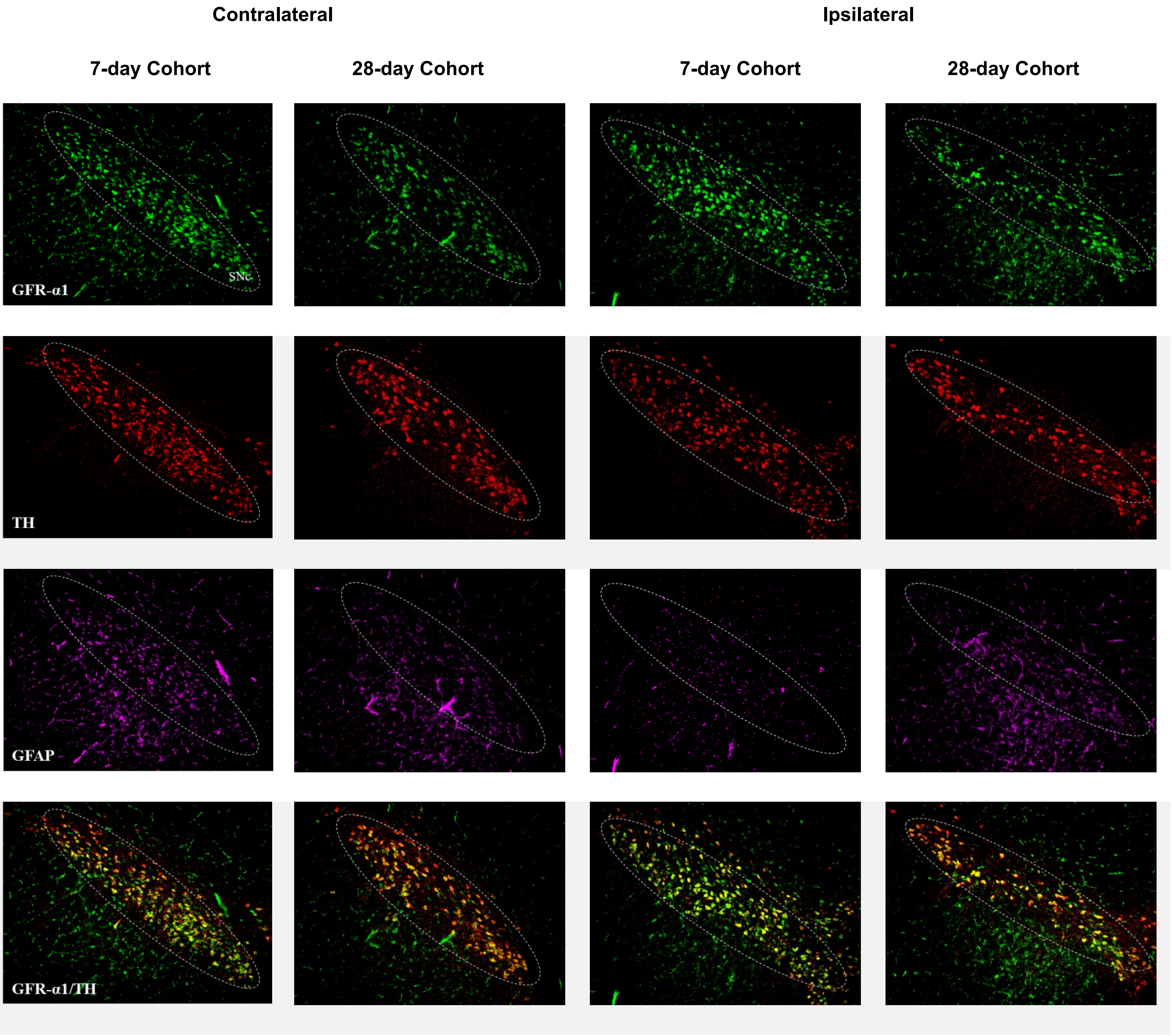

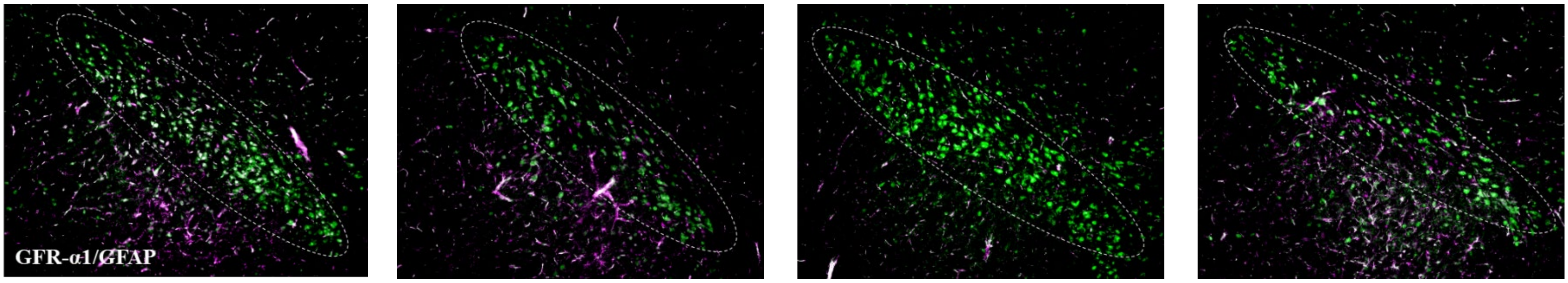


Representative images of GFR-α1 triple staining with TH and GFAP from the contralateral and ipsilateral SN of the sham groups.

**Supplemental Table 1**

| Analyte | region | lesion | days post-Tx | Tx hemisphere |
|---|---|---|---|---|
| **GDNF** | Str | ns, $F_{(1,28)}$=1.17 | ****, $F_{(1,28)}$=34.96 | ns, $F_{(1,21)}$=1.41 |
| **GDNF** | SN | ns, $F_{(1,49)}$=0.95 | ***, $F_{(1,49)}$=13.50 | ns, $F_{(1,49)}$=1.55 |
| **GFAP** | SN | **, $F_{(1,15)}$=9.84 | **, $F_{(1,15)}$=9.43 | ns, $F_{(1,12)}$=0.02 |
| **GFR-α1** | Str | *, $F_{(1,29)}$=7.36 | ns, $F_{(1,29)}$=0.81 | ****, $F_{(1,29)}$=28.5 |
| **GFR-α1** | SN | ***,$F_{(1,50)}$=12.70 | *, $F_{(1,50)}$=6.82 | ns, $F_{(1,20)}$=1.11 |
| **GFR-α1/TH** | SN | ns, $F_{(1,18)}$=2.86 | *, $F_{(1,18)}$=4.47 | **, $F_{(1,17)}$=15.50 |
| **GFR-α1/GFAP** | SN | **, $F_{(1,15)}$=12.68 | **, $F_{(1,15)}$=11.86 | ns, $F_{(1,14)}$=0.66 |
| **RET** | Str | ****, $F_{(1,27)}$=53.0 | ns, $F_{(1,27)}$=1.18 | ****, $F_{(1,20)}$=86.8 |
| **RET** | SN | *, $F_{(1,28)}$=6.82 | *, $F_{(1,28)}$=4.67 | ns, $F_{(1,24)}$=0.38 |
| **BDNF** | Str | ns, $F_{(1,29)}$=0.58 | ns, $F_{(1,29)}$=0.52 | ns, $F_{(1,25)}$=1.50 |
| **BDNF** | SN | ns, $F_{(1,29)}$=0.69 | ns, $F_{(1,29)}$=0.09 | ns, $F_{(1,29)}$=0.14 |
| **TrkB** | Str | ns, $F_{(1,25)}$=2.86 | ns, $F_{(1,25)}$=0.15 | ns, $F_{(1,20)}$=0.68 |
| **TrkB** | SN | ns, $F_{(1,27)}$=3.94 | ns, $F_{(1,27)}$=0.02 | ns, $F_{(1,20)}$=0.68 |

Legend: *$p$<0.05, **$p$<0.01, ***$p$<0.001, ****$p$<0.0001

**Significant interactions:** lesion (L), days post-treatment (DT), treatment hemisphere (TH)

| | | |
|---|---|---|
| **GDNF** | Str | L x TH,****$F_{(1,21)}$=25.70; DT x TH, * $F_{(1,28)}$=4.49 |
| **GDNF** | SN | L x DT x TH, **$F_{(1,49)}$=8.34 |
| **GFR** | Str | L x TH,****$F_{(1,29)}$=30.38; L x DT x TH, $F_{(1,29)}$=4.82 |
| **GFR-α1/TH** | SN | L x TH,**$F_{(1,17)}$=14.73 |
| **RET** | Str | L x TH, ****$F_{(1,20)}$=53.44 |
| **RET** | SN | L x TH, *$F_{(1,24)}$=5.82 |
| **BDNF** | SN | L x TH, **$F_{(1,29)}$=9.61 |

**Statistical analysis results from 3-way ANOVA.** The results from each assay outcome show whether there was an overall effect of 6-OHDA lesion as compared to the sham-operated group (lesion (L)); days past treatment ((Tx) (lesion or sham-op) (DT)); or side of treatment (treatment hemisphere (TH)).
